## Supplementary_materials.zip for "Sensitivity analysis of factors influencing the ecology of mosquitoes involved in the transmission of Rift Valley fever virus": S1_Appendix.pdf

### Calculation of distributions

#### Typical area scanned by *Aedes* flyers

Mosquito dispersal distances are often right-skewed, with most individuals travelling short distances while a few travel much farther. To model this distribution, a Gamma distribution has been chosen, as it is well-suited for positively skewed data and has been used in ecological dispersal studies [1]. The Gamma distribution is parameterized by a shape parameter ($k$) and a scale parameter ($\theta$), which determine the distribution's form. These parameters are estimated using the known mean dispersal distance ($\mu$) and an assumed variance ($\sigma^2$), following the relationships,

$$\mu = k\theta, \quad \sigma^2 = k\theta^2. \tag{1}$$

Given an estimated mean radius distance of 89 m (based on the broadest dataset [2]), the overall expected dispersal distance was calculated as $\mu = \pi \times 89^2$. Dispersal studies have indicated that the distribution is not highly skewed, but it does have a long tail. A lower $k$ (e.g. $k < 1$) would result in a heavily skewed distribution, which may overestimate the proportion of mosquitoes travelling very short distances. On the other hand, a higher $k$ (e.g., $k > 5$) would produce a more symmetric distribution, which may underestimate long-range dispersers. Hence, here we chose $k = 2.5$ to allow for a peak at a common dispersal distance while still maintaining a long right tail. Using this mean and an appropriate choice of $k$, the scale parameter $\theta$ was derived to ensure consistency with empirical data.

#### Number of eggs per batch

We propose that the number of eggs per batch follows a normal distribution. This assumption is supported by several empirical studies cited that report egg counts across different mosquito species and experimental conditions as mean ± standard error or within defined ranges. These distributions generally exhibit symmetric variation around a central mean (e.g., $90 \pm 12$ eggs in *Aedes aegypti* [3]), with no evidence of heavy skew or zero-inflation. While biological counts are non-negative and sometimes skewed, the observed ranges (e.g., 60–125 eggs) do not strongly suggest departure from normality in this case. Hence, in the absence of raw individual-level data, and given the moderate spread and central tendency in reported means and standard deviations, the normal distribution provides a reasonable and tractable approximation for modeling fecundity.

To estimate the parameters of this normal distribution, specifically the mean ($\mu$) and standard deviation ($\sigma$), we combine the available data from the various studies. To estimate the overall mean ($\mu_{\text{overall}}$), we calculate the arithmetic mean of the means reported in each study. This provides a central estimate of the number of eggs per batch across all studies, and is computed as

$$\mu_{\text{overall}} = \frac{\sum_{i=1}^{n} \overline{X_i}}{n}, \tag{2}$$

where $\overline{X_i}$ is the mean from study $i$ and $n$ is the number of studies. If a mean is provided as a range, we use the midpoint of the range as the best estimate for $\overline{X_i}$.

To estimate the overall standard deviation ($\sigma_{\text{overall}}$), we account for the variability in the means across the studies. The overall standard deviation is estimated by computing the root-mean-square of the standard deviations (or standard errors) from the studies, which captures the spread of all the data across all studies. The formula for the overall standard deviation is,

$$\sigma_{\text{overall}} = \sqrt{\frac{\sum_{i=1}^{n} \sigma_i^2}{n}}, \quad (3)$$

where $\sigma_i$ is the standard deviation reported in study $i$. This approach provides a reasonable estimate of the spread of the data across the studies.
