## Supplementary_materials.zip for "Sensitivity analysis of factors influencing the ecology of mosquitoes involved in the transmission of Rift Valley fever virus": S1_Table.pdf

**Table S1. Summary of literature search for the typical area scanned by flyers. The citation number is linked to the main manuscript.**

| Citation | Mosquito | Measure/keywords | Estimation |
|---|---|---|---|
| [26] | *Aedes vexans* | Linear regression, | 1394 m (Fig. 3) |
| | *Culex neavei* | maximum distance | 1356 m (Fig. 3) |
| | *Culex poicilipes* | | 843 m (Fig. 3) |
| [27] | *Cx. quinquefasciatus* | Isotype enrichment, dispersal, | 59% $1-2$ km from natal larval habitat |
| | | seeking oviposition | 15% $>$ 2 km (Fig. 10B) |
| | | | 26% $<$ 1 km (Fig. 10B) |
| | *Ae. albopictus* | Host seeking | 100% $<$ 1 km |
| | | | 79% $<$ 0.25 km |
| [28] | *Cx. Pipiens* | Mean distance travelled (MDT) | Total, min: 0.16 km, max: 1.98 km |
| | *Cx. salinarius* | | MDT: 1.33 km (Table 1) |
| [29] | *Ae. lineatopennis* | Mean distance travelled post emergence | 0.15 km (Table 2) |
| [30] | *Ae. aegypti* | Mean distance travelled, weighted mean distance | 106 m (95% CI: 87.68, 123.69) (Fig. 2) |
| [31] | *Ae. aegypti* | Oviposition traps, mean | 50% $<$ 32 m |
| | | dispersal distance, | mean 45.2 m (95% CI: 39.7, 51.3) |
| | | Laplacian kernel | 10% $>$ 100 m |
| [32] | *Culex* | Average flight distance, | Avg. 609.5 m (SD 437.0 m) (Table 4) |
| | | average maximum distance | Max. 5014 m (Table 3) |
| | *Aedes* | | Avg. 89.0 m (SD 50.1 m) |
| | | | Max. 2959 m (Table 3) |
