## Supplementary_materials.zip for "Sensitivity analysis of factors influencing the ecology of mosquitoes involved in the transmission of Rift Valley fever virus": S2_Appendix.pdf

### Parameters

The values for each parameter that have been used throughout the stability analysis, which can be found in the supplementary material of [1]. For ease and reproducibility, they have been presented in Table S2.

environment if table fits in text column.

**Table S2. Parameter values used when running the code.**

| Parameter | Symbol in Code | Value |
|---|---|---|
| *Change for Stability Analysis* | | |
| Number of days in simulation | day_length | $365 * 11$ |
| Constant Temperature (Table 1) | constanttemp | 25 |
| Constant Water Body Area (Table 1) | constantwb | 17 500 |
| Livestock Total | livestocktotal | 500 |
| Detection probability, $p_f$ (Table 1) | prop_find | 0.1 |
| *Remain for Total Analysis* | | |
| Time Step | delta_t | 1 |
| Number of ponds in each cell | N_ponds | 1 |
| Reduction rate for *Culex*, based on Figure 8 in [2] | reduction_C | 0.5 |
| Reduction rate for *Aedes*, based on Figure 8 in [2] | reduction_A | 0.58 |
| Birth rate for livestock | b_L1 | 1/(5 * 365) |
| Death rate for livestock | mu_L1 | 1/(5 * 365) |
| Parameter for the impact of the livestock on vector fecundity and gonotrophic cycles | q_divided | 1.00E+11 |
| Probability of transovarial transmission | q_A | 0.007 |
| Latent Period | epsilon_L1 | 2/7 |
| Infectious Period | gamma_L1 | 1/30 |
| Average time for egg deposition | t_dep | 0.229 |
| *Parameters for Periodic Functions* | | |
| Frequencies of oscillations in surface areas of water bodies | omega_S_p | 2*3.14/(365) |
| Frequencies of oscillations in surface areas of temperature | omega_T_a | 2*3.14/(365) |
| Mean surface area of water bodies during periods $2\pi/\omega_S$ | C_S_p | 17500 |
| Mean temperature during periods $2\pi/\omega_T$ | C_T_a | 21 |
| The maximum amplitudes in the water oscillations | A_S_p | (1-0.4)*C_S_p |
| The maximum amplitudes in the temperature oscillations | A_T_a | 7 |
| Respective water phases | phi_S_p | acos(-1)† |
| Respective temperature phases | phi_T_a | acos(-1)† |

†Values for phases are equal to this when the periodic functions are in phase (i.e. reach a peak at the same time). We use phi_T_a = acos(0) for out of phase.

Note that the daily larva mortality and daily pupa mortality for *Culex* are defined, respectively, as

$$\mu_L^{Culex} = 37.9317808331 - 0.2573339304 \cdot T + 0.0004364566 \cdot T^2 \tag{1}$$

$$\mu_P^{Culex} = 80.3113158804 - 0.5439116495 \cdot T + 0.0009210259 \cdot T^2, \tag{2}$$

where $T$ is the temperature measured in Kelvins. Similarly, the daily larva and pupa mortality for *Aedes* are defined, respectively, as

$$\mu_L^{Aedes} = 50.1205 - 0.3393650263 \cdot T + 0.0005747698 \cdot T^2 \tag{3}$$

$$\mu_P^{Aedes} = 3.524873 - 0.023943082 \cdot T + 0.00004066735 \cdot T^2. \tag{4}$$

If the rounded figures are used from the supplementary material [1], then a negative death rate occurs. These are defined as stated here in the code.
