## Supplementary_materials.zip for "Sensitivity analysis of factors influencing the ecology of mosquitoes involved in the transmission of Rift Valley fever virus": S3_Appendix.pdf

### Investigation into parameters

To better understand the changes observed in the Sobol indices, we conducted a targeted investigation into how the development rates, mortality rate, and oviposition rate affect the model when temperature and water body area are held constant (Figs. S4–S6).

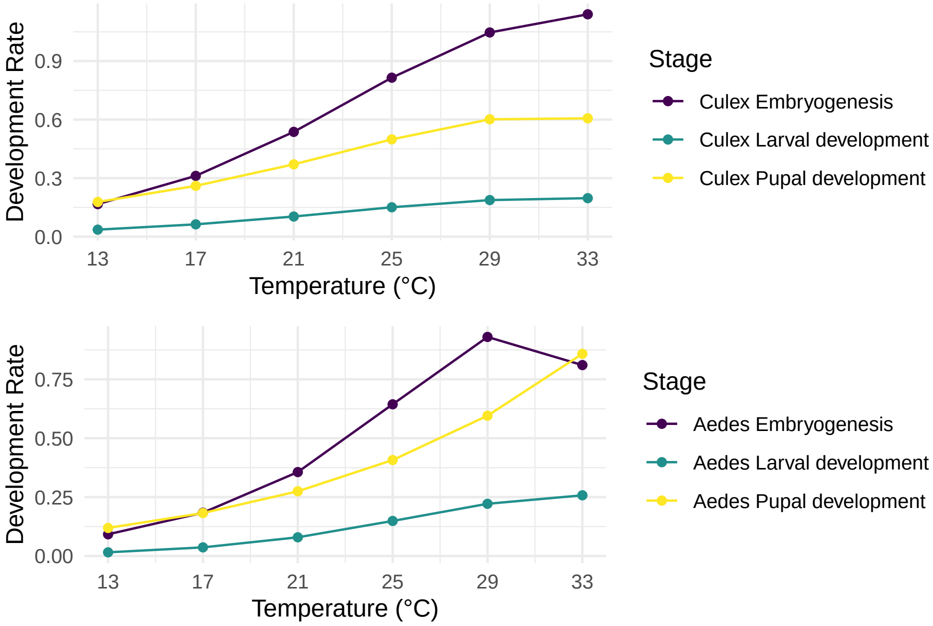


**Fig S4. Comparison of development rates for different constant temperature values.** For A: *Culex*. B: *Aedes*. Increases in development rates are observed as temperature increases.

Fig S4 highlights that as temperature increases, then so does the development rate. However, the order of rate does not change for the temperature range that we are investigating. That is, the rate from egg to larvae remains the highest, followed by the rate from pupae to adult, and the lowest rate is from larvae to pupae, suggesting that numerous eggs convert to larvae before the other stages catch up (since the larvae are moving to pupae slowly). Whilst the pupae do leave faster than they enter into the compartment, there may not be enough supply.

Fig S5 highlights that there are little changes in the mortality rate between the chosen temperatures (21-29 degrees). There is a marginal increase in the mortality rate for *Culex* pupae, and *Aedes* larvae. A notable mortality rate are the culex eggs, which is over 0.75, with a considerable jump between 13-17 degrees and 29-33 degrees.

Finally, the oviposition rates are displayed in Fig S6. The oviposition rates depend on the water body area. It can be observed that increases in the water body area increases the oviposition rate.

Taken together, these results provide useful context for interpreting the patterns observed in the Sobol sensitivity analysis. By examining development, mortality, and oviposition rates under constant environmental conditions, we gain a clearer

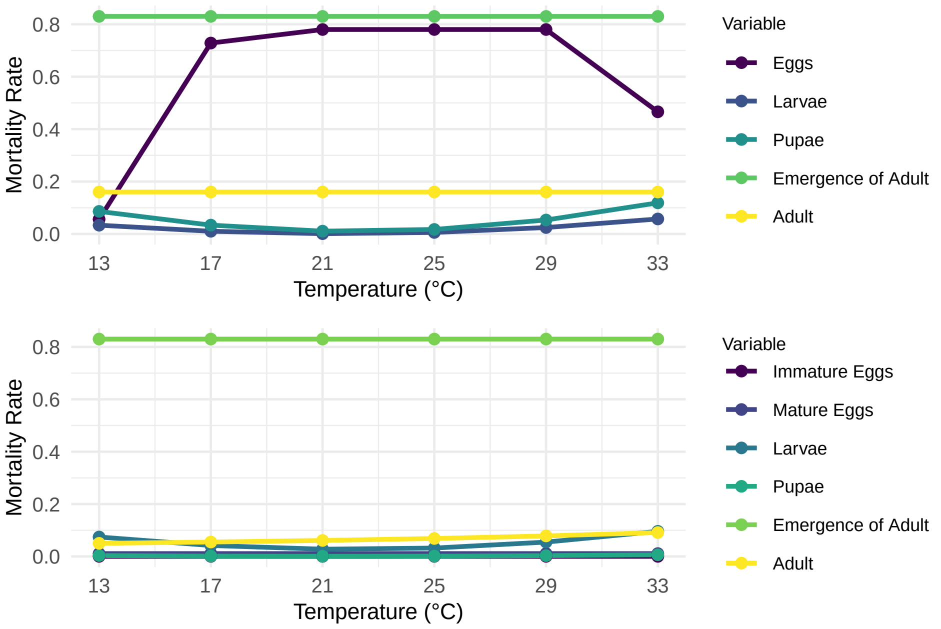


**Fig S5. Comparison of mortality rate for the different constant temperature values.** For A: *Culex*. B: *Aedes*. There is very little change between the rates. There is no dependence on the size of the water body area.

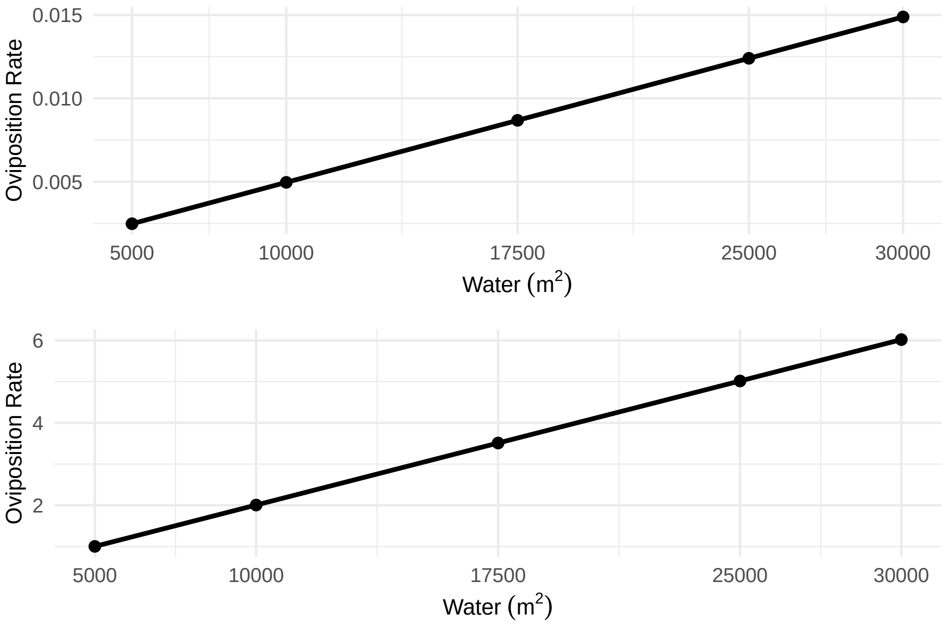


**Fig S6. Comparison of oviposition rate for different constant temperature values.** For A: *Culex*. B: *Aedes*. A linear change is seen as water bodies increases.

understanding of how individual parameters behave within the model. The trends observed, such as the consistent ordering of development rates, limited variation in mortality across temperatures, and the dependence of oviposition on water body area—are broadly consistent with findings in the literature. While not definitive, this targeted exploration offers a useful reference point for interpreting the relative influence of these parameters in the model.
