## Supplementary_materials.zip for "Sensitivity analysis of factors influencing the ecology of mosquitoes involved in the transmission of Rift Valley fever virus": S3_Fig.pdf

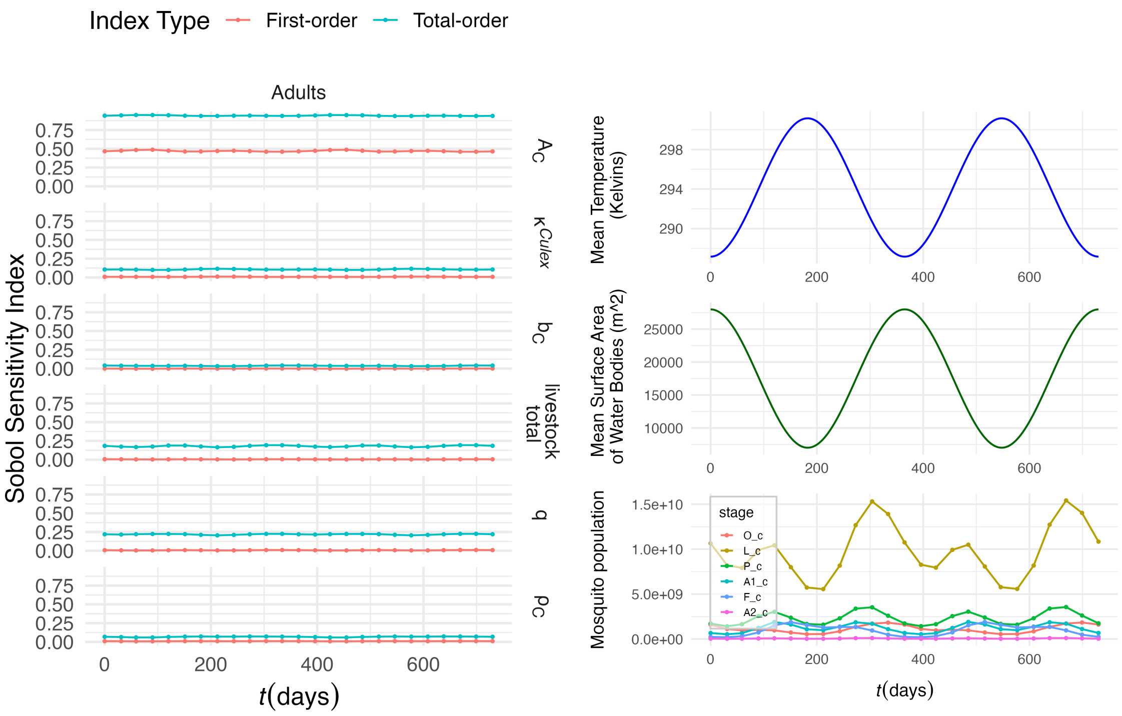


**(a)** *Culex*

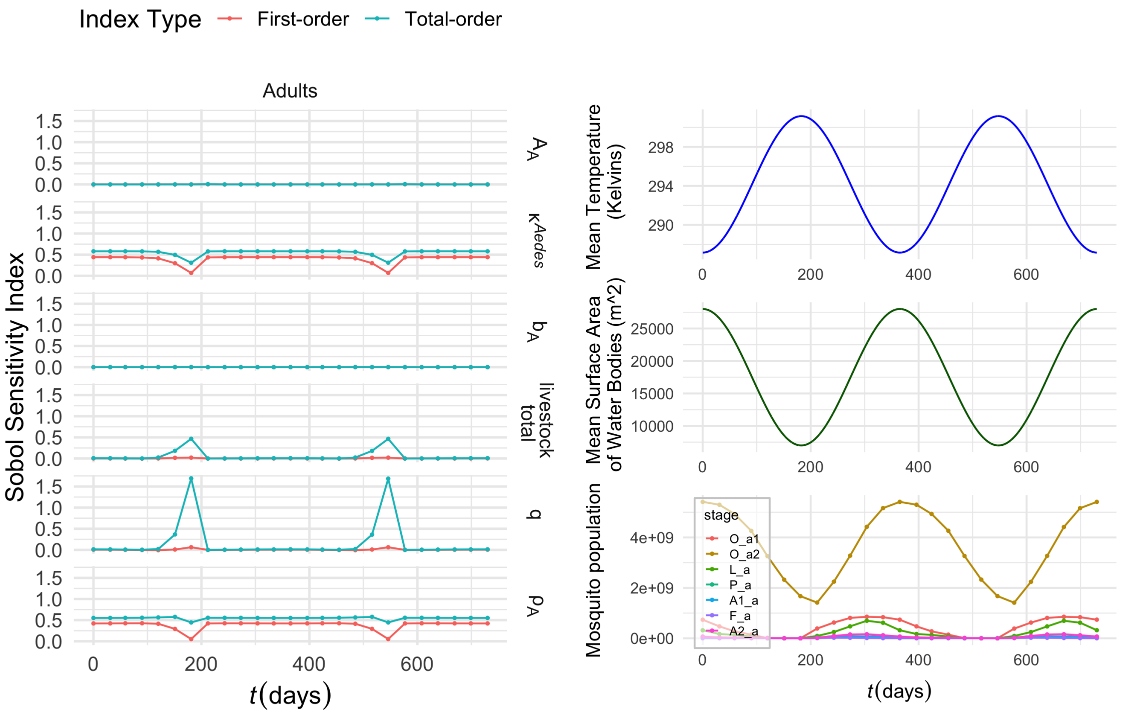


**(b)** *Aedes*

**Fig S3.** Time varying Sobol sensitivity indices for *Culex* and *Aedes* with out-phase periodic functions for temperature and water bodies.
