## Supplementary_materials.zip for "Sensitivity analysis of factors influencing the ecology of mosquitoes involved in the transmission of Rift Valley fever virus": S4_Appendix.pdf

#### Second order Sobol indices

Within this appendix, we investigate Sobol indices to a higher degree, looking at second order indices. Higher order indices are used to describe the fraction of the output variance due to interactions between uncertain input parameters [1]. Second order indices are denoted as $S_{ij}$ and describes the fraction of the output variance due to interaction between input parameters $i$ and $j$. Due to computational cost, this investigation is conducted only for the medium, constants scenario.

##### Method

Many systems exhibit non-linear relationships, in which the influence of individual inputs cannot be fully understood without also considering how they act together. In such cases, two inputs may have minimal effect when varied independently, but their combined variation can produce a substantial change in the model output. To quantify these joint effects, we compute second-order Sobol sensitivity indices, which explicitly measure the contribution of pairwise interactions to the total output variance. Mathematically, these indices are defined as

$$S_{ij} = \frac{V[E[Y \mid X_i, X_j]] - V[E[Y \mid X_i]] - V[E[Y \mid X_j]]}{V[Y]}, \tag{1}$$

where $Y$ is the output, and $X_i$, $X_j$ are the input parameters of interest. Calculating the second order indices allows us to identify synergistic or compensatory relationships that cannot be captured through first-order indices alone.

Whilst, it can be sufficient to conduct a Sobol sensitivity analysis for first and total order indices with $N(m+1)$ model evaluations [2], Saltelli [3] recommends using $N(2m+2)$ evaluations to obtain more reliable results. Here, $N$ denotes the Monte Carlo sample size, and $m$ is the number of input parameters in the model. This requirement highlights that more extensive sampling is needed when accounting for higher-order effects, which correspondingly increases the computational cost of the analysis.

For this analysis, the R package *sensobol* [4] was used to extend the original analysis and compute the second- and third- order Sobol indices, with the sampling matrices configured by specifying the order.

##### Results

The second-order Sobol sensitivity analysis revealed that only a small subset of parameter pairs contributed meaningfully to the *Culex* model's output variance (Fig. S7). The majority of the interactions displayed indices close to zero, indicating limited non-additive effects among the majority of the parameters. In contrast, stronger interactions emerge for pairs involving $q$ and livestock total, where when they are paired with the area scanned the index number is approximately 0.09. Although these were the highest interaction effects detected, a value of this magnitude still represents a small proportion of the total variance, and therefore does not indicate a strong interaction in absolute terms.

It is important to note that the relatively large second-order indices associated with the area scanned by *Culex* is likely to reflect substantial knowledge uncertainty, highlighting the need to measure this factor more robustly.

On the other hand, the second-order analysis for *Aedes* revealed only a single pair contributed meaningfully to the model's output variance (Fig. S8). This pair corresponded to the same

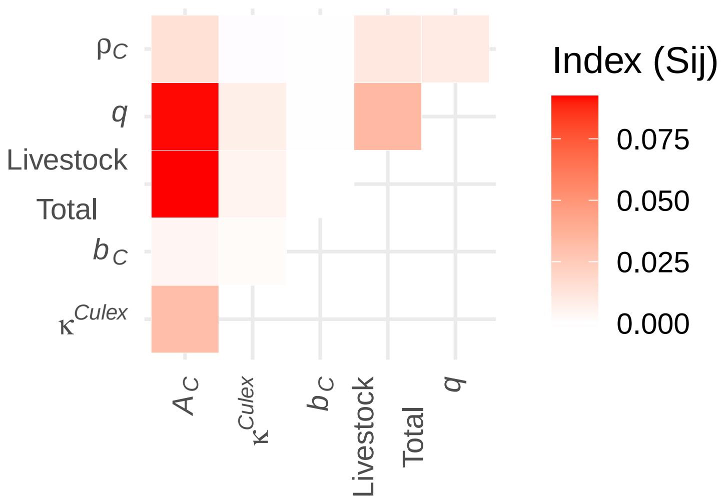


**Fig S7. Second order Sobol sensitivity indices for *Culex*.** Matching the analysis for the baseline first and total order indices with constant temperature and water bodies.

parameters identified as influential in the first- and total-order analyses: the maximum egg density and the proportion of soil area suitable for oviposition. The alignment between first-order, total-order, and second-order results indicates that variation in these two parameters, both individually and in combination, is the primary driver of uncertainty in the *Aedes* model.

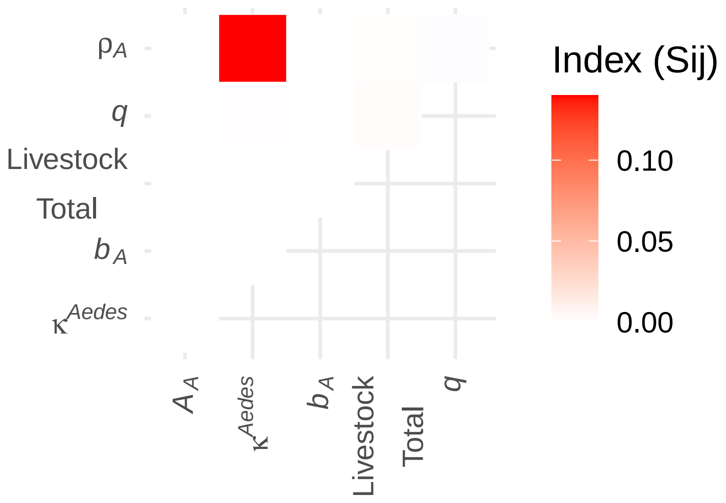


**Fig S8. Second order Sobol sensitivity indices for *Aedes*.** Matching the analysis for the baseline first and total order indices with constant temperature and water bodies.

### References

1. Zhang XY, Trame MN, Lesko LJ, Schmidt S. Sobol Sensitivity Analysis: A Tool to Guide the Development and Evaluation of Systems Pharmacology Models. CPT: Pharmacometrics &

Systems Pharmacology. 2015;4(2):69. doi:10.1002/PSP4.6.

2. Tetsuya Homma, Homma T, Andrea Saltelli, Saltelli A. Importance measures in global sensitivity analysis of nonlinear models. Reliability Engineering & System Safety. 1996;52(1):1–17. doi:10.1016/0951-8320(96)00002-6.

3. Saltelli A, Tarantola S, Campolongo F, Ratto M. Sensitivity Analysis in Practice: A Guide to Assessing Scientific Models. Wiley; 2004.

4. Puy A, Piano SL, Saltelli A, Levin SA. sensobol: An R Package to Compute Variance-Based Sensitivity Indices. Journal of Statistical Software. 2022;102(5):1–37. doi:10.18637/jss.v102.i05.
