## Supplementary_materials.zip for "Sensitivity analysis of factors influencing the ecology of mosquitoes involved in the transmission of Rift Valley fever virus": S5_Appendix.pdf

### Investigation into differences

The Sobol matrices are based on the distributions observed in Table 3. The minimum and maximum values for each parameter can be seen in Table S3, along with the range and the percentage deviation from the midpoint. A higher percentage indicates a wider spread of values relative to the midpoint. If the percentage is low, it means the range is small compared to the midpoint, implying less variability.

**Table S3. Parameter range in Sobol Matrix for $N = 1500$ samples.**

| Parameter | Minimum | Maximum | Range | Percent |
|---|---|---|---|---|
| $\mathcal{A}_C$ | 464 866 | 15 593 548 | 15 128 682 | 188% |
| $\mathcal{A}_A$ | 783 | 102 848 | 102 065 | 197% |
| $\kappa^{Culex}$ | 0.001 48 | 0.9985 | 0.997 02 | 199% |
| $\kappa^{Aedes}$ | 0.001 48 | 0.9985 | 0.997 02 | 199% |
| $b_C$ | 95.6 | 202.3 | 106.7 | 72% |
| $b_A$ | 45.7 | 64.2 | 18.5 | 34% |
| $\rho_C$ | 519 043 | 19 990 479 | 19 471 436 | 190% |
| $\rho_A$ | 20 957 | 999 521 | 978 564 | 192% |

Notably, the parameter representing the number of eggs laid per batch has the smallest percentage range. Combining insights from the literature review and this sensitivity analysis, it becomes evident that this parameter is the least influential compared to the three other parameters under investigation. This suggests that selecting a value within its given range results in minimal impact on the overall outcomes.

To examine the impacts of the parameter ranges on the models, we ran the sensitivity analysis within the *Culex* model with the *Aedes* parameters and the *Aedes* model with the *Culex* parameters (Fig S9). It can be observed that the order of significance of the Sobol indices have remained the same, with minimal differences between the size of the index.

Thus, the consistency in the order of Sobol indices across models, despite the swapped parameter ranges, indicates that the observed differences are primarily driven by the extent of parameter variability rather than any structural or functional issues within the models themselves.

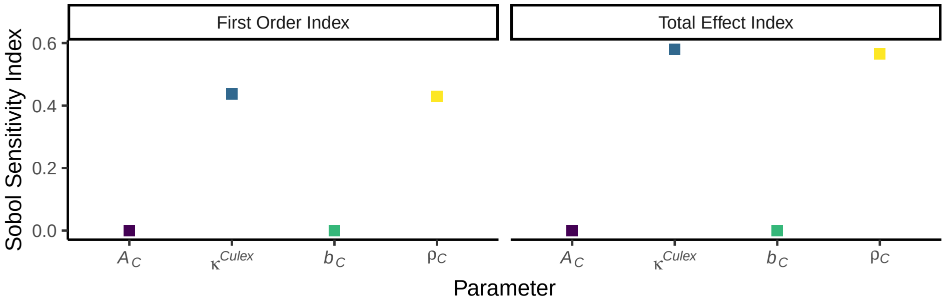


A

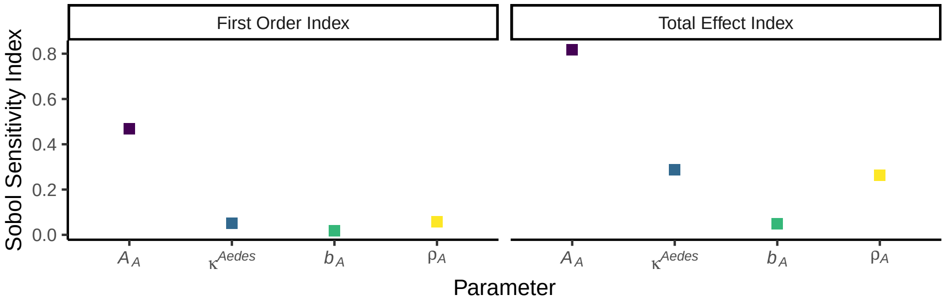


B

**Fig S9. First order and total effect Sobol sensitivity indices with medium values of temperature, water body area and detection probability**. A: *Culex* mosquitoes using the *Aedes* parameter ranges for the sensitive parameters. B: *Aedes* mosquitoes using the *Culex* parameter ranges for the sensitive parameters.
