## Supplementary figures and images for "Sensitivity analysis of factors influencing the ecology of mosquitoes involved in the transmission of Rift Valley fever virus"

### S1_Fig.pdf

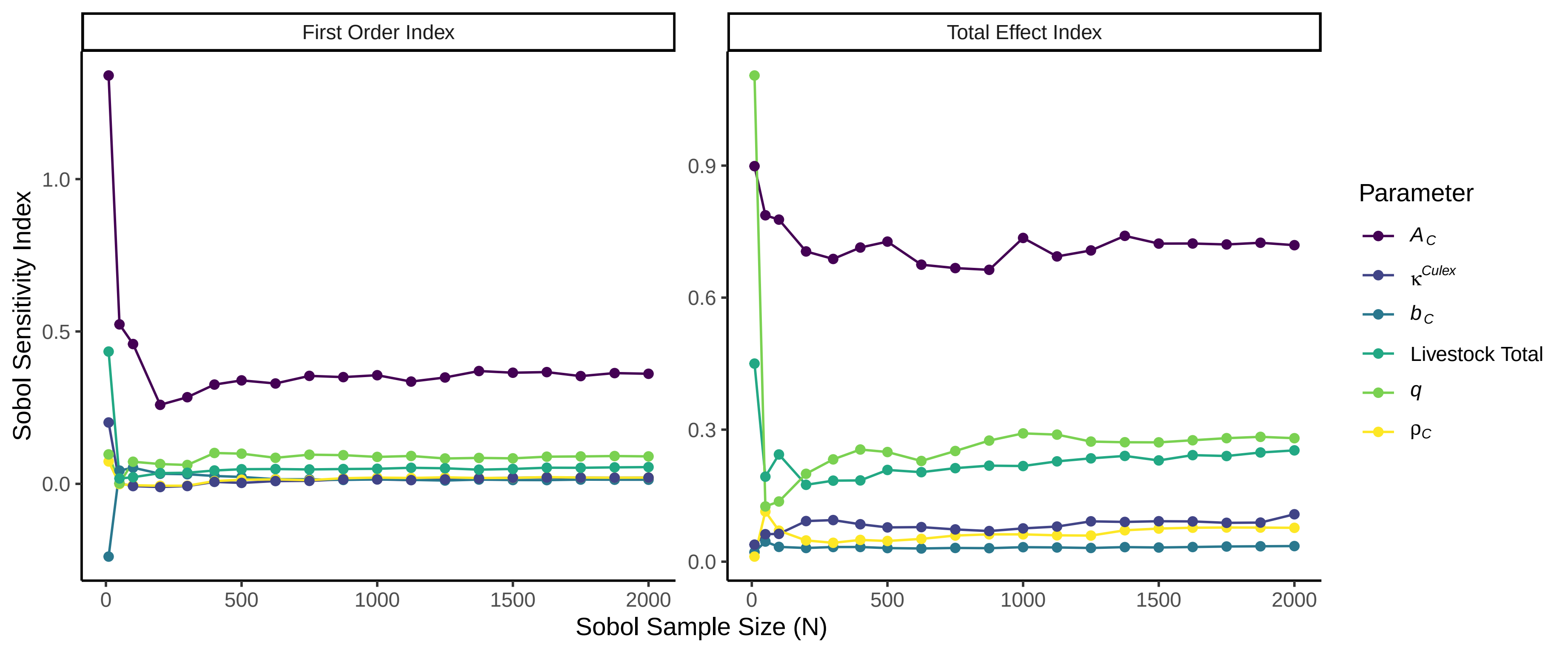

### S2_Fig.pdf

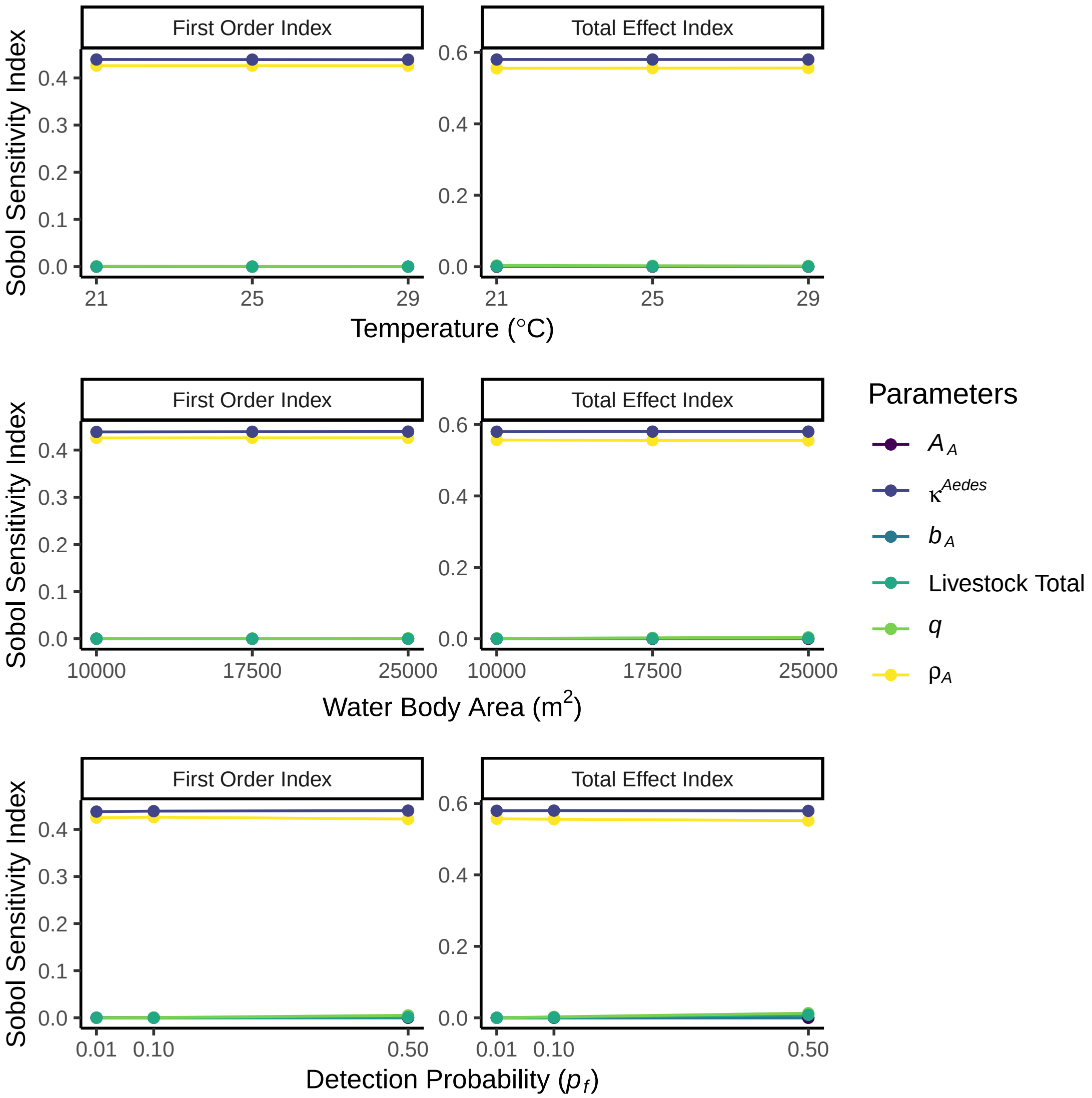
